# You Only Look Once (YOLO) Based Machine Learning Algorithm for Real-Time Detection of Loop-Mediated Isothermal Amplification (LAMP) Diagnostics

**DOI:** 10.1101/2025.03.21.644632

**Authors:** Biniyam Mezgebo, Ryan Chaffee, L. Ricardo Castellanos, S. Ashraf, J. Burke-Gaffney, Johann D. D. Pitout, Bogdan I. Iorga, M. Ethan MacDonald, Dylan R. Pillai

## Abstract

Loop-mediated isothermal amplification (LAMP) is a widely used rapid and affordable molecular DNA amplification method with minimal resource requirements. However, visual interpretation of results is subjective and prone to errors, leading to potential false-positive and negative results. To address this limitation, a machine-learning approach is proposed for automated LAMP classification based on digital images. The approach utilizes You Only Look Once (YOLOv8), a fast and robust object detection algorithm to locate and classify tubes within LAMP images, enabling automated categorization as positive or negative. The trained model achieved a high overall accuracy of 95.5% in classifying LAMP images into positive or negative. Additionally, the approach had a 98.0% precision and 92.7% recall for positive cases and 93.4% precision and 98.2% recall for negative cases, demonstrating its potential for real-time LAMP diagnosis and enhanced assay performance. This project demonstrated the platform’s suitability for real-time testing, offering an easy operation and rapid results.

## Introduction

Diagnosing infectious diseases like malaria and COVID-19 relies on various techniques, each with limitations. Traditional methods, including pathogen culture, microscopy, and PCR-based diagnostics, often face cost, resource, and turnaround time limitations. While Quantitative Polymerase Chain Reaction (qPCR) offers high accuracy and is considered the gold standard for specific tests, its limitations include the need for specialized equipment, skilled personnel, and complex protocols, hindering its use in point-of-care (POC) settings (1,2). In this context, isothermal loop-mediated amplification (LAMP) is a compelling alternative for POC diagnostics. LAMP’s simplicity and ability to amplify minimally processed or raw samples make it particularly suitable for resource-limited settings (3,4). These advantages have fueled LAMP’s exploration in various fields, especially POC diagnostics for pathogens like SARS-CoV-2 (5). Several POC platforms have been developed, demonstrating LAMP’s potential for rapid and affordable detection (6–9). However, LAMP typically relies on qualitative visual detection, which can introduce ambiguity and potentially lead to misdiagnosis. This work proposes to address this limitation by using machine learning approaches with LAMP, aiming to automate result interpretation while maintaining LAMP’s POC advantages.

Integrating deep learning-based detection and prediction methods with existing diagnostic tools is gaining momentum, aiming to produce more time-efficient results and automate interpretation, ultimately assisting healthcare professionals in making informed data-driven decisions (10). Automated diagnostics can also minimize human-related errors, benefiting patients and healthcare systems. In recent years, researchers have proposed diverse techniques for developing machine learning-driven portable and low-cost LAMP devices for rapid pathogen detection. These approaches range from simple rule-based systems to traditional machine learning (ML) and deep learning (DL) network-based methodologies (10,11). In the context of LAMP signal analysis, researchers have explored a wide spectrum of ML algorithms to develop efficient classifier models. These include convolutional neural networks (CNNs), linear discriminant analysis, random forests, gradient boosting classifiers (GBCs), Bayesian networks, fuzzy logic, and decision trees (10,11). For example, Rohaim *et al*. demonstrated the efficacy of a CNN model in identifying different colours in LAMP images (10). Similarly, Song *et al*. developed a machine learning-based smartphone application to analyze SARS-CoV-2 diagnostics results using a LAMP with clustered regularly interspaced short palindromic repeats (CRISPR), referred to as the DAMPR assay system (11). In their work, ML techniques were employed to quantify the SARS-CoV-2 gene concentration by analyzing 300 sample images for each concentration. The study compared LDA, RF, and GBC classifiers, with RF achieving the best accuracy of 99.38%.

YOLO, introduced by Redmon *et al*. (2016) (12), is a real-time object detection algorithm that leverages convolutional neural networks (12–14). Unlike conventional two-stage detectors such as R-CNN and Fast R-CNN, which first generate region proposals (RPs) potentially containing objects and then classify them using CNNs, YOLO adopts a single-stage approach (15–18). It divides the input image into a grid and simultaneously predicts bounding boxes, object presence, and class probabilities for each cell, streamlining the detection process. This unified approach significantly reduces processing time compared to multi-stage methods (19–26).

YOLO has different versions; among recent advancements, YOLOv5 and YOLOv7 are the most popular variants. YOLOv5 leverages deep learning methodologies and a Cross-Stage Partial (CSP) network structure to improve efficiency and overall performance (26). However, it still struggles with small object detection and challenging scenarios. YOLOv7, on the other hand, introduced the Trainable Bag of Freebies (TBoF) strategy, enhancing accuracy and generalization through data augmentation and other techniques. While effective, YOLOv7 can be sensitive to training data and model parameters, and it often demands substantial computational resources (27).

This paper focuses on applying YOLOv8, the latest iteration of the YOLO model, for LAMP image analysis. The YOLOv8 builds upon the strengths of its predecessors while introducing several key improvements: YOLOv8 surpasses previous YOLO versions in terms of accuracy, particularly in challenging scenarios involving small objects or dense environments. This enhanced precision makes it a more reliable tool for LAMP detection, leading to potentially more accurate diagnoses. Also, YOLOv8 strongly emphasizes real-time object detection, making it highly suitable for practical applications requiring rapid results. This characteristic is crucial for efficient diagnosis of malaria, COVID/RSV, and antimicrobial resistance (AMR) among others, in field settings, where timely diagnosis can significantly impact patient outcomes.

In this study, we harness YOLOv8’s efficiency and accuracy to automate the interpretation of LAMP results. By formulating the classification of LAMP images as a multi-object detection problem, we develop a model capable of precisely locating tubes within LAMP images and classifying them as “Positive” or “Negative” based on visual characteristics. This integration of machine learning with LAMP technology offers several advantages: it streamlines the diagnostic process, reduces human error, and enables real-time disease diagnosis. Moreover, the speed of YOLO aligns well with LAMP’s rapid amplification, potentially creating a synergistic effect that could significantly accelerate the overall testing process. This study demonstrates the potential of combining advanced object detection algorithms with isothermal amplification techniques to enhance the efficiency and reliability of molecular diagnostics.

## Results

This section discusses the results of applying yolov8 to LAMP images. Fig 1 shows the prediction results of the proposed research. The positive samples are marked with red bounding boxes, while the negative samples are highlighted with green bounding boxes. Each detected sample in the images is assigned a confidence score/probability that reflects the model’s certainty that a given detection corresponds to a real object and its corresponding class. A confidence threshold above 0.5 was utilized in the algorithm to determine the validity of the detections and their classifications.

**Fig 1.**
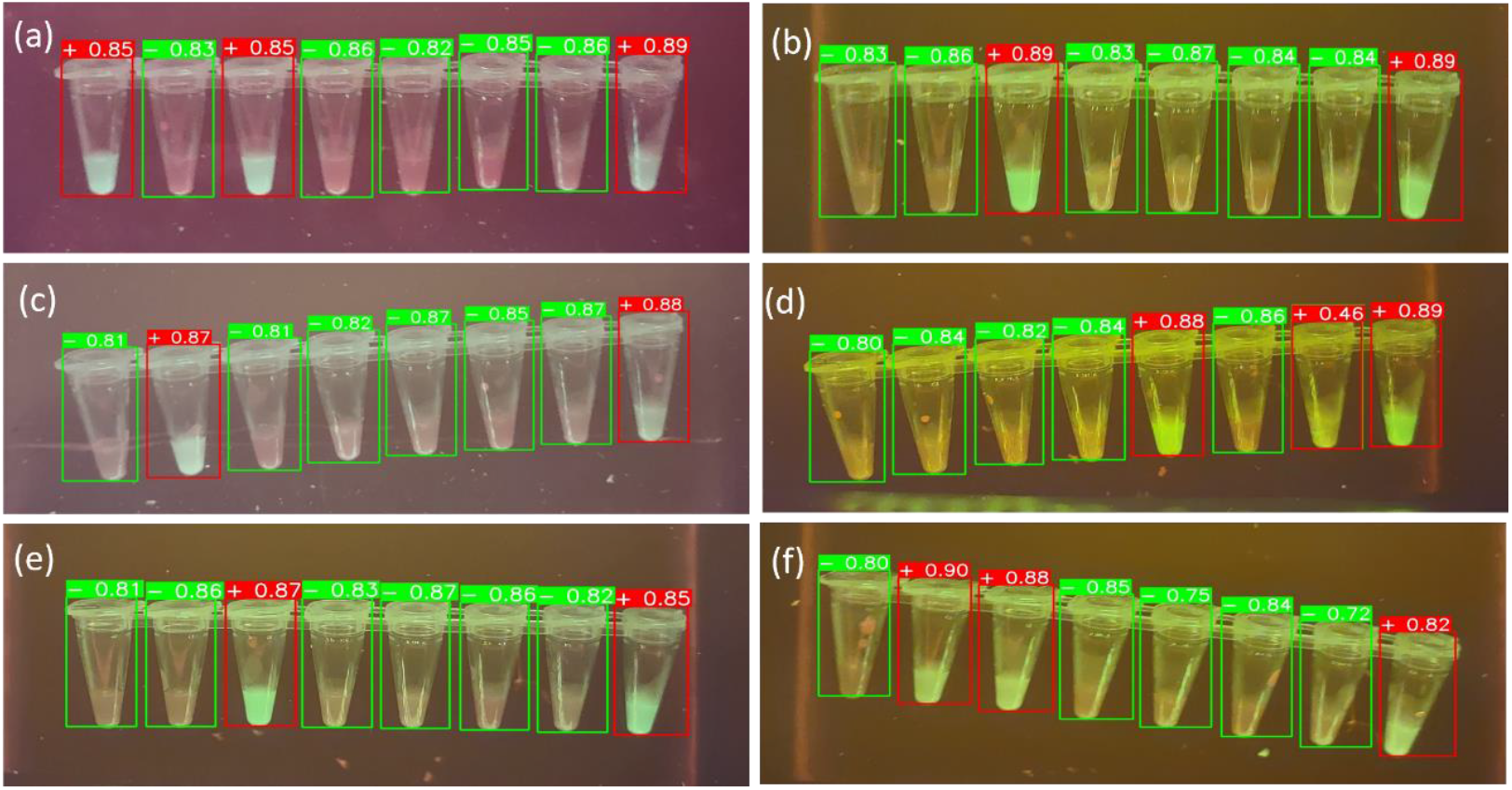
Example images of the predictions result of the trained model on unseen data. (a) - (f) example images from the “white pearl” demonstrate the capacity of the algorithm to detect and classify positive and negative results of individual wells in 8-tube LAMP strips. The model demonstrates robust detection performance across different illumination conditions and color variations. Green and red bounding boxes indicate negative and positive LAMP reactions respectively, with confidence scores shown above each detection. The model maintains high detection accuracy (confidence scores ranging from 0.72 to 0.96) despite challenging variations in: (1) background illumination (purple to brown tones), (2) fluorescence intensity (bright to dim), (3) sample opacity, and (4) image capture angles.

Fig 2 shows the loss values for the box loss, object loss, and class loss at each epoch during the training and validation process. The box loss represents the difference between the predicted and ground-truth bounding box coordinates, the object loss represents the confidence score for each object detected in an image, and the class loss represents the probability of each detected object belonging to a specific class. Training an object detection model aims to minimize the total loss, combining box, object, and class loss. The loss values exhibit a decreasing trend as the training progresses, indicating an improvement in the model’s ability to detect tube samples in the images.

**Fig 2.**
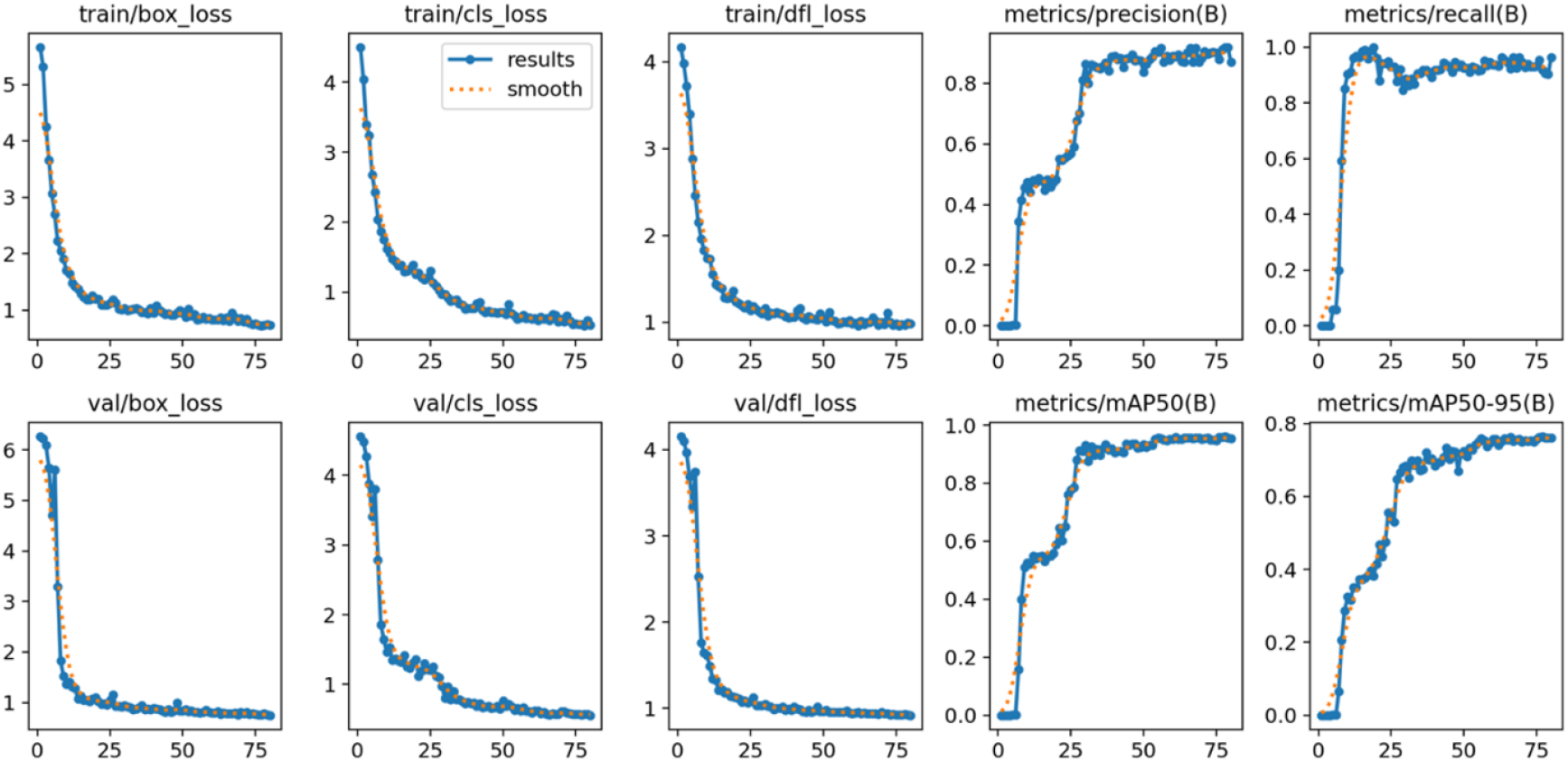
Training and validation metrics for YOLOv8 model on LAMP image detection. The plots show progression of various loss functions (box, class, and dfl losses) for both training and validation sets for 80 epochs. Performance metrics including precision, recall, mAP@50, and mAP@50-95 are also displayed, demonstrating the model’s improving accuracy in detecting and classifying LAMP assay results. The smooth convergence of loss functions and the upward trends in accuracy metrices indicate successful training and good generalization of the model for LAMP image analysis.

The trained YOLOv8 model achieved promising performance in classifying LAMP reaction images, as shown in Fig 1. The model was evaluated on a validation set of 45 images containing 566 instances of tube images (Table 1), where it demonstrated good overall accuracy and individual class performance. The model achieved a mAP@50 of 0.97 and a mAP@50-95 of 0.79, indicating high precision in detecting and classifying LAMP images across various intersection-over-union thresholds. This signifies exceptional accuracy in both identifying and localizing LAMP images. The model’s performance was particularly strong in distinguishing between positive and negative classifications, with mAP@50 values of 0.988 and 0.97 respectively. Furthermore, the model demonstrated high overall precision (0.91) and recall (0.97) on the validation set, reinforcing its effectiveness in LAMP image analysis.

**Table 1.** Performance metrics of the YOLOv8 model on validation data for LAMP Image classification. The model was evaluated on 45 validation images containing 566 instances across two classes (positive and negative). Overall performance and class-specific metrices are reported, including Precision, Recall, mAP@50, and mAP@50-95.

| Class | Images | Instances | Precision | Recall | mAP@50 | mAP@50-95 |
| --- | --- | --- | --- | --- | --- | --- |
| all | 45 | 566 | 0.915 | 0.978 | 0.976 | 0.799 |
| Positive |  | 260 | 0.927 | 0.981 | 0.988 | 0.811 |
| Negative |  | 306 | 0.983 | 0.934 | 0.971 | 0.786 |

Fig 3 shows the confusion matrix for our model. It reveals strong overall performance with 286 true negatives and 255 true positives out of 566 total instances. The model achieved an accuracy of 95.5% with a precision of 98.1% for positive class detection and a recall of 92.7%. The model exhibited excellent specificity, correctly identifying 98.2% of negative samples. While the false positive rate was low (5 instances), there were slightly more false negatives (20 instances), suggesting a conservative bias in positive detection. This bias may be preferable in diagnostic contexts where false positives would lead to more serious consequences than false negatives.

**Fig 3.**
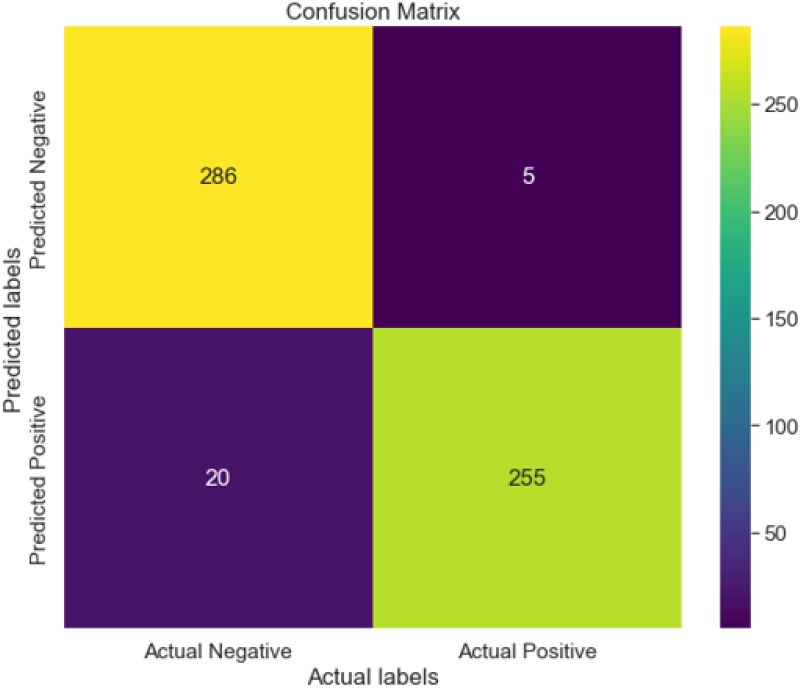
Confusion matrix demonstrating high-accuracy classification of LAMP images.

Fig 4 presents a comprehensive set of evaluation metrics that complement the results shown in Table 1, providing a deeper analysis of our model’s performance. The figure includes precision-confidence (a), recall-confidence (b), precision-recall (c), and F1-confidence (d) curves for both individual classes and overall performance. The precision-recall curve (c) demonstrates the model’s ability to maintain high precision across a wide range of recall values, with <u>mAP@0.5</u> reaching 0.959 for all classes. This indicates robust performance in balancing false positives and false negatives. The F1-confidence curve (d) shows a peak F1 score of 0.92 at a confidence threshold of 0.696, suggesting an optimal balance between precision and recall at this point. Furthermore, the precision-confidence and recall-confidence curves (a, b) illustrate the model’s high confidence in its predictions, maintaining high precision and recall values across a range of confidence thresholds.

**Fig 4.**
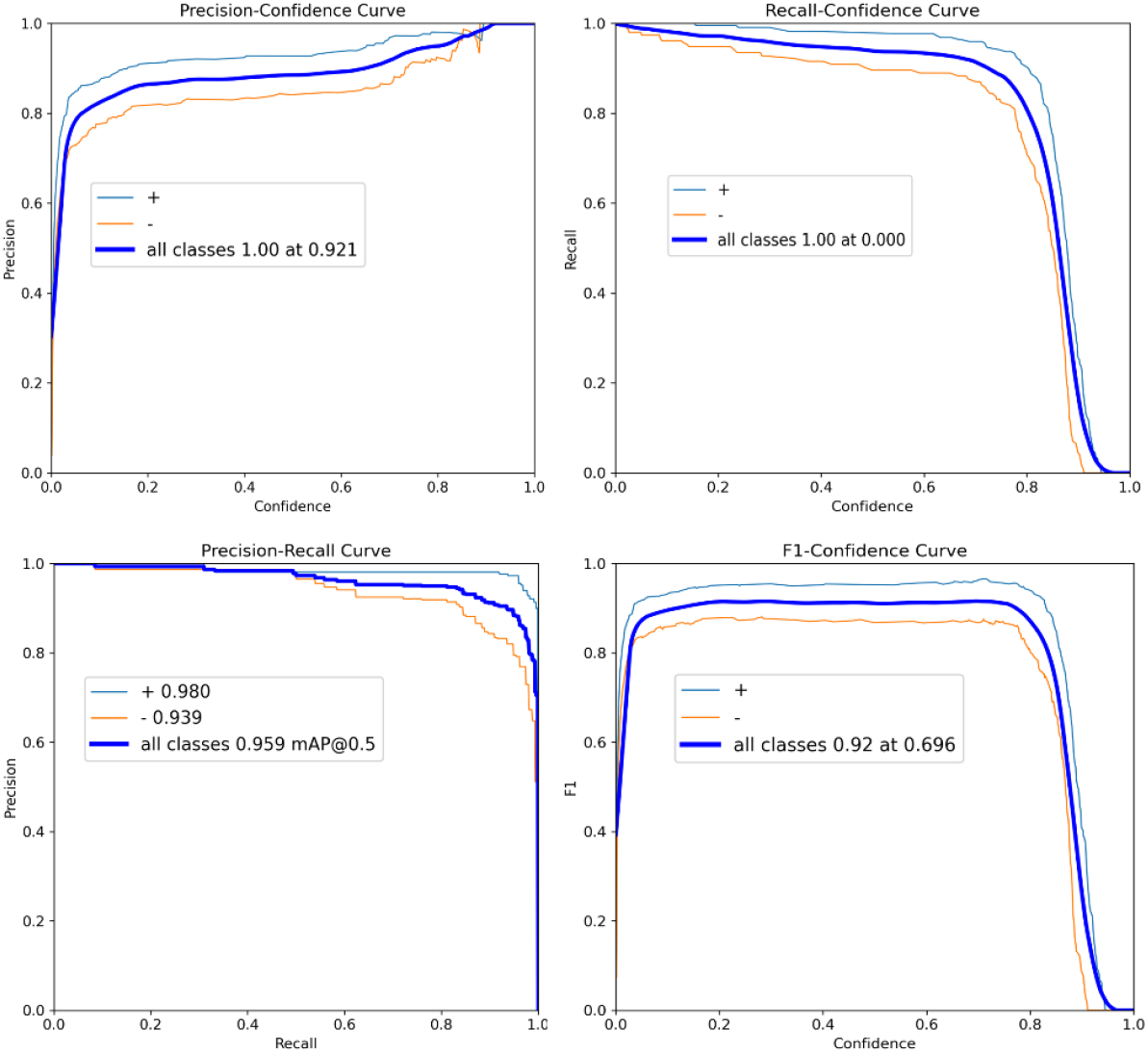
Performance Metrics. (a) Precision-Confidence, (b) Recall-Confidence, (c) Precision-Recall, (d) F1-Confidence

The high precision and recall values, in addition to the confusion matrix results, indicate that the model successfully minimized both false positives and false negatives, which is crucial for diagnostic applications. The strong performance of the model across different illumination conditions and color variations, as evidenced by the example images, suggests good generalization capabilities. This is particularly important for LAMP-based diagnostics, where lighting conditions and sample characteristics may vary between laboratories and point-of-care settings. The confidence scores shown in the detection results indicate that the model not only makes correct classifications, but also has high certainty, which is essential for reliable automated analysis.

While the model achieved promising results, this study had several limitations and areas for future improvement. The slightly higher number of false negatives compared to false positives suggests that the model could benefit from additional fine-tuning to capture subtle positive signals better. Furthermore, while the model performed well on the current dataset, future work should evaluate its robustness on a more diverse set of LAMP images, through the use of different imaging devices and diagnostic environments.

## Materials and Methods

### YOLOv8 Algorithm and Architecture

This sub-section provides a brief description of YOLOv8, one of the fastest and most accurate object detection algorithms (28). The YOLO model architecture is typically structured into three primary components: the backbone, neck, and head, as shown in Fig 5.

**Fig 5.**
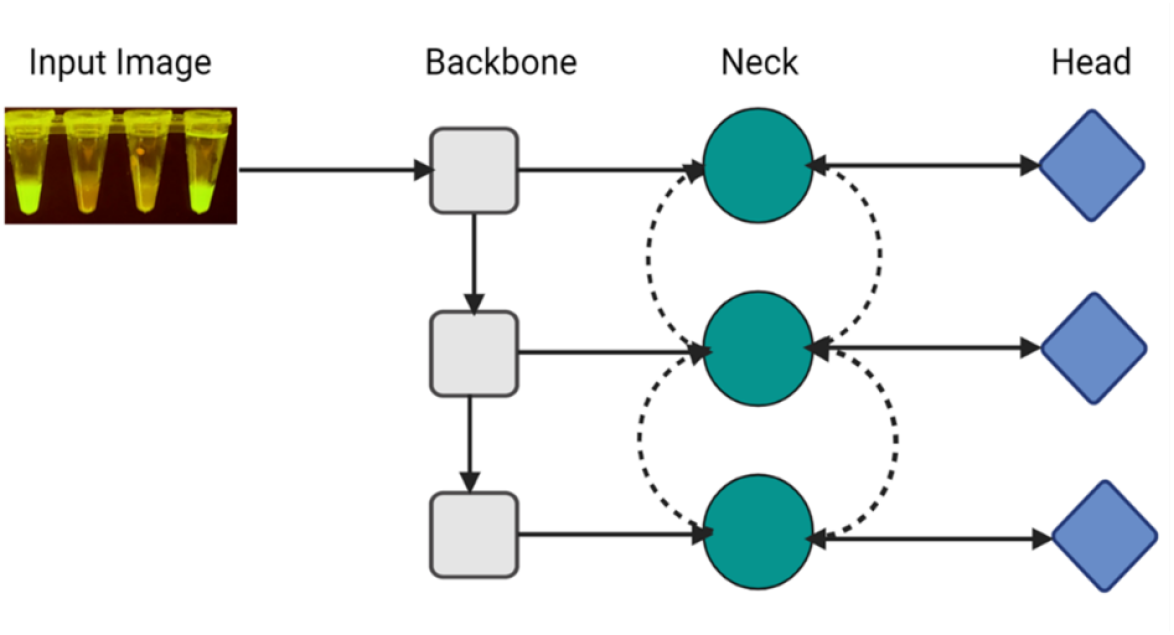
The architecture of YOLO. The input image shows a LAMP test result with fluorescent reactions. The YOLO architecture consists of three main components: (a) the backbone for hierarchical feature extraction at different scales, capturing both fine and coarse details, (b) the neck (green circles with bidirectional dotted arrows) for multi-scale feature aggregation and enhancement. The bidirectional arrows indicate feature fusion and information flow between different scales allowing the model to better understand multi-scale features. (c) the head (blue diamonds) for final predictions: object classification and bounding box regression. Having three heads suggest that predictions are made at three different scales which is typical in YOLO architectures for detecting objects of various sizes. The three rows represent different feature map scales for detecting various object sizes: top(small), middle (medium), and bottom (large).

The architectural design incorporates an initial input layer that receives an image and a series of convolutional layers that systematically extract relevant information from the image. Each component is crucial in feature extraction, processing, and final output generation.

- **Backbone**: consists of convolutional layers that extract features from the input images at varying scales. This component, often a pre-trained CNN like VGG16 or ResNet50, extracts valuable features from the input image. Lower-level features and high-level features are extracted on the shallow and deeper layers.
- **Neck**: The neck is an intermediate component connecting the backbone and head. It is responsible for combining feature maps from different backbone layers using modules like Path Aggregation Networks (PAN) or Feature Pyramid Network (FPN).
- **Head**: This component is responsible for final predictions; the head generates bounding boxes, objectness scores, and class labels.

Fig. 6 shows the model architecture of YOLOv8. Continuous improvements in the three components have significantly enhanced the overall accuracy and speed of the YOLO network. YOLOv8 further revolutionized object detection by adopting an anchor-free approach and refining the grid-based prediction system. It divides the input image into an *M*×*M* grid, where each grid cell directly predicts objects. For each detected object, YOLOv8 predicts N values: *x, y, w, h, θ, c*_1_ ⋯ *c*_*K*_. Where, (*x, y*) represents the center coordinates of the object relative to the grid cell, (*w, h*) denote the width and height of the object relative to the entire image, *θ* is the objectness score indicating the model’s confidence in the presence of an object, and (*c*_1_, *c*_2_, ⋯, *c*_*K*_) are the class probabilities for K different classes.

**Fig 6.**
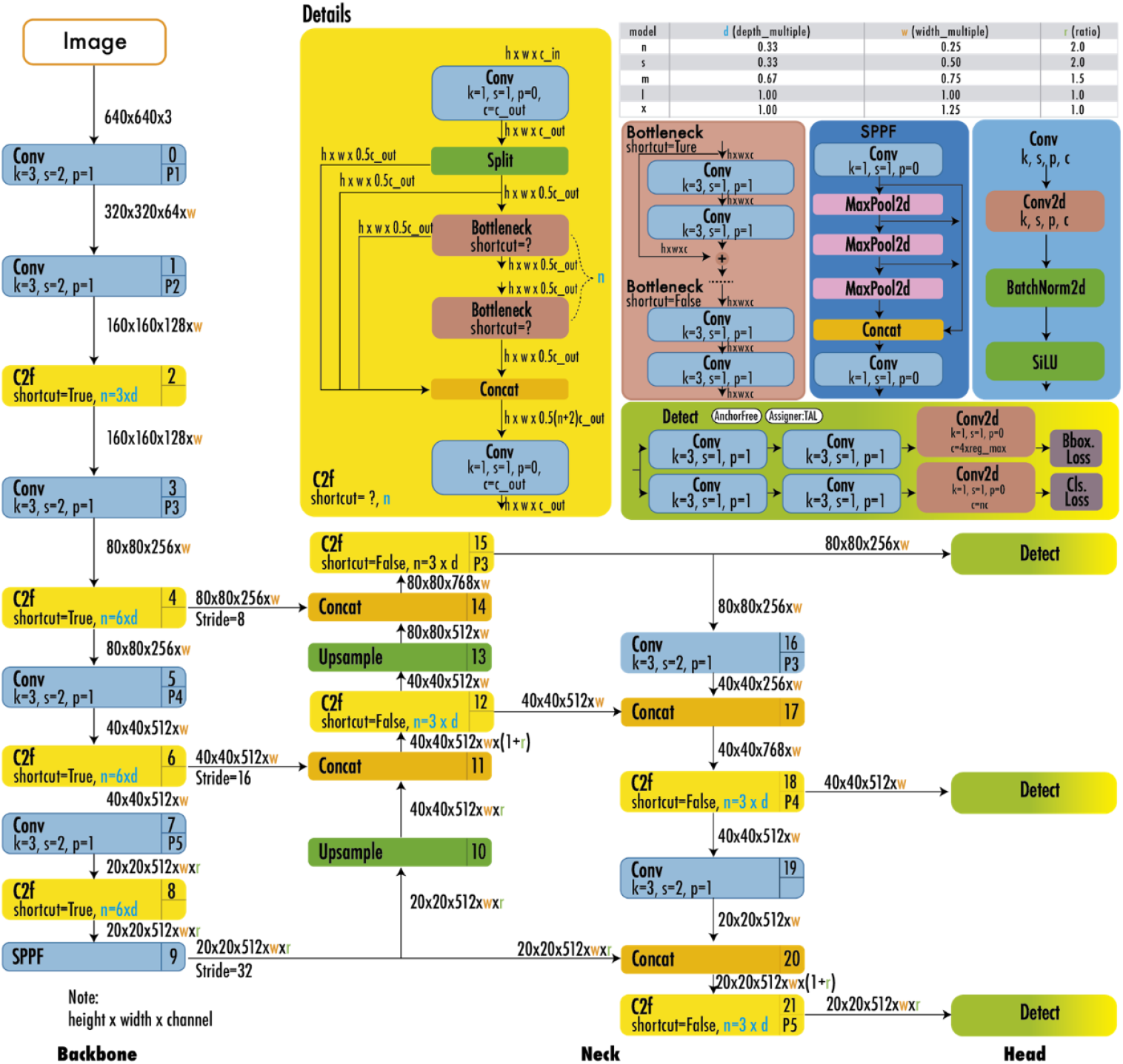
The architecture of YOLOv8 consists of a backbone, neck, and head, adopted from (28).

**Fig 7.**
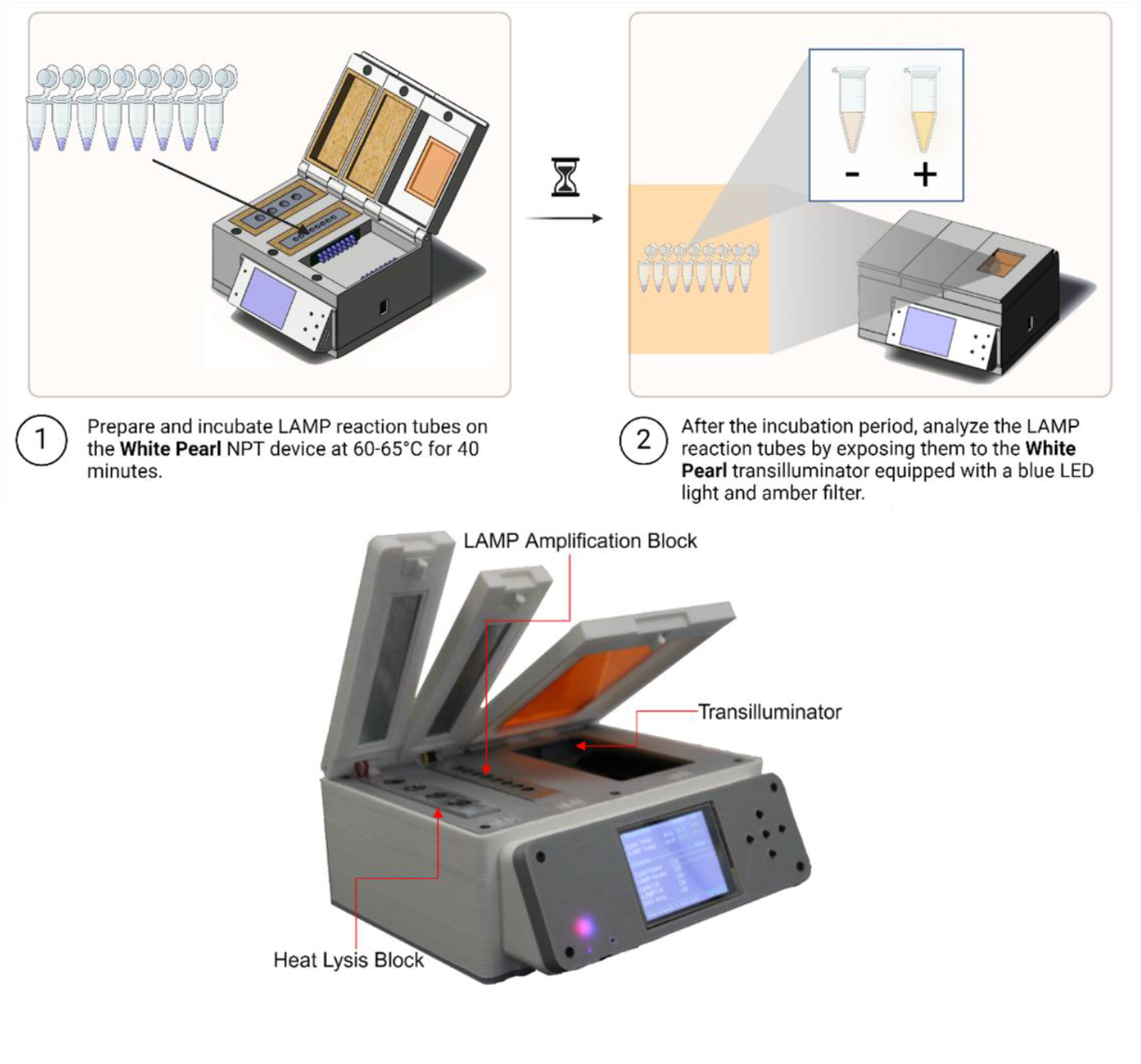
White Pearl LAMP device.

The output of YOLOv8 is a tensor of size *H*×*W*×(4 + 1 + *K*), where *H*×*W* is the dimensions of the feature maps at each prediction scale, 4 represents the bounding box coordinates, 1 is for the objectness score, and K is the number of classes. YOLOv8 uses the maximum class probability as an indicator of object presence. The model uses a sigmoid function in the output layer as the activation function for class probabilities, representing the likelihood of an object belonging to each possible class (28). Further, the output of YOLOv8 undergoes post-processing steps to refine predictions: A confidence threshold is applied to filter out low-confidence predictions and non-maximum suppression (NMS) is used to eliminate redundant detections. YOLOv8’s grid cells efficiently handle tasks related to object localization and classification. The model estimates the probability of an object’s center falling within each grid cell, as formulated by Equation 1:

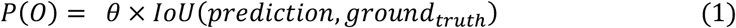

where: *P*(*O*) is the detection confidence for object *O, θ* is the objectness score, and *IoU (prediction, ground_truth)* is the Intersection over Union between the predicted and actual bounding boxes and quantifies the overlap between predicted and ground truth bounding boxes. YOLOv8 uses *IoU* to determine acceptable detection areas and to make decisions about object localization.

### Experimental Setup and Model Training

As a proof of principle, LAMP tests designed to detect antimicrobial resistance genes in bacteria were utilized. Samples consisted of a variety of gram-negative bacteria with extended spectrum beta-lactamase and carbapenemase genes. These tests were performed as previously described (29). The tests were performed using an in-house developed device dubbed the White Pearl (Fig 3). This device consisted of three primary modules – (i) Thermal Lysis module: A dedicated heating block maintains a constant temperature of 95°C for initial sample lysis, (ii) Isothermal Amplification module: A separate heating block ensures consistent isothermal amplification at 64°C and (iii) Fluorescence Detection module: A transilluminator equipped with two 2×16 side-mounted blue light emitting diode (LED) arrays, and an absorptive filter of orange acrylic.

In this work, 3612 LAMP tube sample images were captured using various Android and iOS smartphones under varied lighting conditions to reflect real world scenarios. Sample images were captured on the white pearl transilluminator and a commercially available transilluminator (examples are shown in Fig 8). This approach is crucial to train a robust model that can handle diverse image conditions, including colour variations, varied capture angles and inconsistent lighting typically encountered when using smartphone cameras.

**Fig 8.**
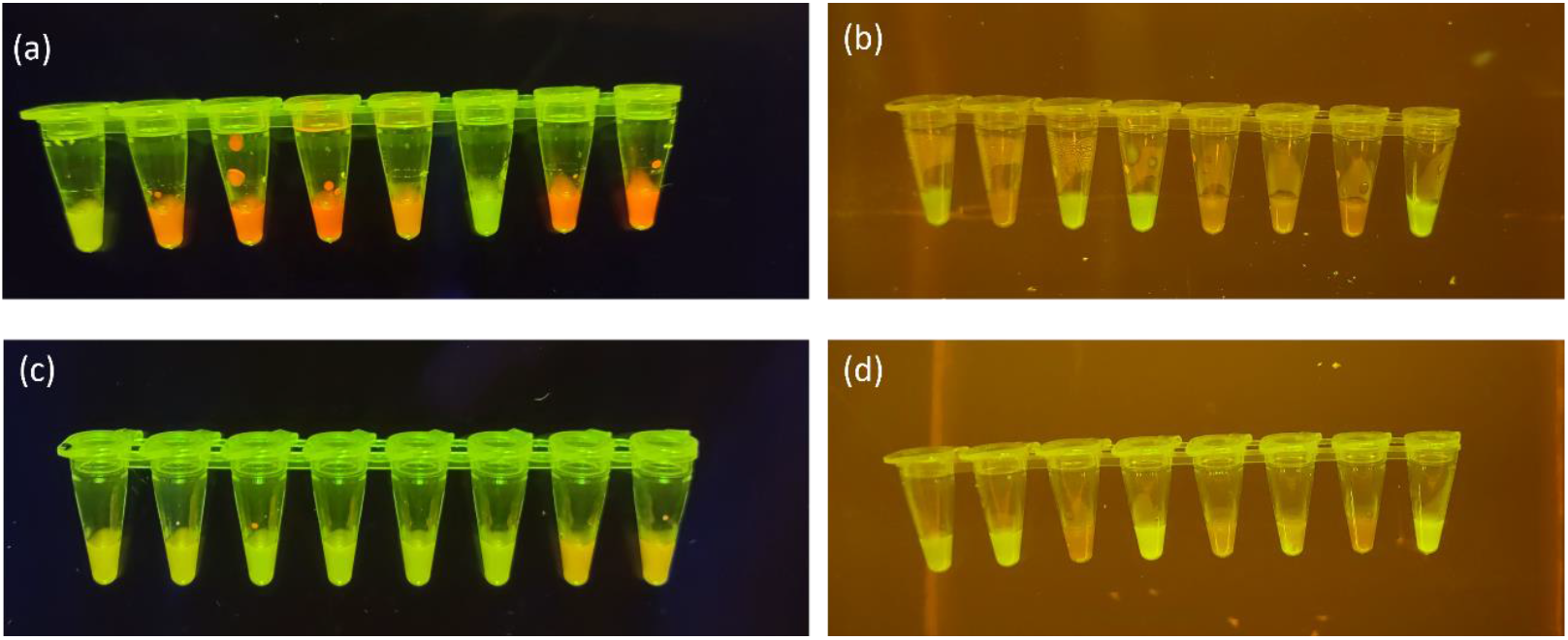
Training dataset examples for YOLOv8 algorithm to detect and classify LAMP assay results. Images depict 8-tube LAMP strips with positive results fluorescing bright green and negative results as a dull orange. (a) and (c)are images captured on a commercial transilluminator. (b) and (d) are images captured using the white pearl transilluminator.

Manual annotation was performed on each image using the Computer Vision Annotation Tool (CVAT), where bounding boxes were drawn to delineate individual samples. The image labels were subsequently saved in .txt format. Positive and negative classifications were assigned based on bacterial isolates’ whole genome sequencing (WGS) results.

Furthermore, before model training, the dataset underwent preprocessing steps, including Min-Max normalization to standardize pixel intensities. This preprocessing step is essential for optimizing model convergence, reducing sensitivity to lighting variations, and enhancing numerical stability in the YOLOv8 model. Following the preprocessing steps, the data was randomly split into training (84%, 3046 images) and validation (16%, 566 images) sets to facilitate model training and evaluation. Also, the training set comprised 1462 positive (48%) and 1584 negative (52%) images and the validation set contained 260 positive (46%) and 306 negative (54%) images.

The model was trained on a machine with an NVIDIA RTX 3060 12GB GPU, Intel i7-10700F CPU operating at 2.90 GHz, 64 GB of RAM, and a Windows 10 64-bit operating system. Python version 3.11 and the PyTorch deep learning framework were used for script execution. The training process utilized images resized to 640 × 640, a batch size of 8, a learning rate 0.01, and 80 epochs.

### Evaluation Metrics

To validate our model’s performance, we used several established evaluation metrics commonly used in object detection tasks. These metrics, including precision, recall, and F1 score, were used to assess the performance of the trained YOLOv8 interms of bounding box quality and class label predictions (30). We used the Precision-Recall curve, a graphical representation of the trade-offs between precision and recall at varied thresholds to visualize the model’s performance. The Average Precision (AP) which computes the area under this curve, provided a single value to indicate the model’s precision and recall performance. For our multi-object detection scenario, we extended this concept to the Mean Average Precision (mAP), calculating the average AP values across multiple object classes to gain broader insight into the model’s effectiveness. We specifically focused on mAP@50, which calculates mAP at an intersection-over-union (*IoU*) threshold of 0.50. mAP50 gives more insight into the model’s accuracy for easier-to-detect objects. The *IoU* metric, measuring the overlap between the model’s predicted bounding box and the ground truth (correct) bounding box, served as a key indicator of localization accuracy. Furthermore, we used mAP@50-95, averaging mAP values across *IoU* thresholds from 0.50 to 0.95, to provide a more comprehensive view of the model’s performance across varying degrees of localization precision. This multi-faceted evaluation approach allowed us to thoroughly assess our YOLOv8 model’s capabilities in detecting and classifying LAMP assay results.

## Conclusions

In conclusion, this study demonstrates the successful application of the YOLOv8 algorithm for automated classification of LAMP reaction results. By leveraging convolutional neural networks and single-pass image processing, our model achieved an impressive overall accuracy of 95.5% in distinguishing between positive and negative samples. The high precision (98.0% for positive, 93.4% for negative) and recall (92.7% for positive, 98.2% for negative) values, along with balanced F1-scores exceeding 95% for both classes, underscore the robustness of our approach. Our methodology, which incorporated a diverse dataset of smartphone-captured images from both Android and iOS devices, proves the model’s adaptability to real-world conditions. This adaptability, combined with the model’s ability to process images in real-time on edge devices, positions our solution as a valuable tool for point-of-care diagnostics. By automating the interpretation of LAMP results, we have addressed the limitations of manual visual interpretation, potentially enhancing the scalability and reliability of LAMP-based molecular techniques.

The integration of deep learning with LAMP technology not only improves assay performance but also opens new avenues for rapid, accurate, and accessible disease diagnosis. As we continue to refine and expand this approach, it holds promise for transforming point-of-care testing practices, particularly in resource-limited settings where rapid and reliable diagnostics are crucial.

## Notes

### Competing Interest Statement

The authors have declared no competing interest.

